# The CD28-transmembrane domain mediates chimeric antigen receptor heterodimerization with CD28

**DOI:** 10.1101/2020.09.18.296913

**Authors:** Yannick D. Muller, Duy P. Nguyen, Leonardo M.R. Ferreira, Patrick Ho, Caroline Raffin, Roxxana Beltran Valencia, Zion Congrave-Wilson, Theodore Roth, Justin Eyquem, Frederic Van Gool, Alexander Marson, James A. Wells, Jeffrey A. Bluestone, Qizhi Tang

## Abstract

Anti-CD19 chimeric antigen receptor (CD19-CAR)-engineered T cells are approved therapeutics for malignancies. The impact of the hinge (HD) and transmembrane (TMD) domains between the extracellular antigen-targeting and the intracellular signaling modalities of CARs has not been systemically studied. Here, a series of CD19-CARs differing only by their HD (CD8/CD28/IgG_4_) and TMD (CD8/CD28) was generated. CARs containing a CD28-TMD, but not a CD8-TMD, formed heterodimers with the endogenous CD28 in human T cells, as shown by co-immunoprecipitation and CAR-dependent proliferation to anti-CD28 stimulation. This dimerization depended on polar amino-acids in the CD28-TMD. CD28-CAR heterodimerization was more efficient in CARs containing a CD8-HD or CD28-HD as compared to an IgG_4_-HD. CD28-CAR heterodimers did not respond to CD80 and CD86 stimulation but led to a significant reduction of CD28 cell-surface expression. These data unveil a new property of the CD28-TMD and suggest that TMDs can modulate CAR T-cell activities by engaging endogenous partners.

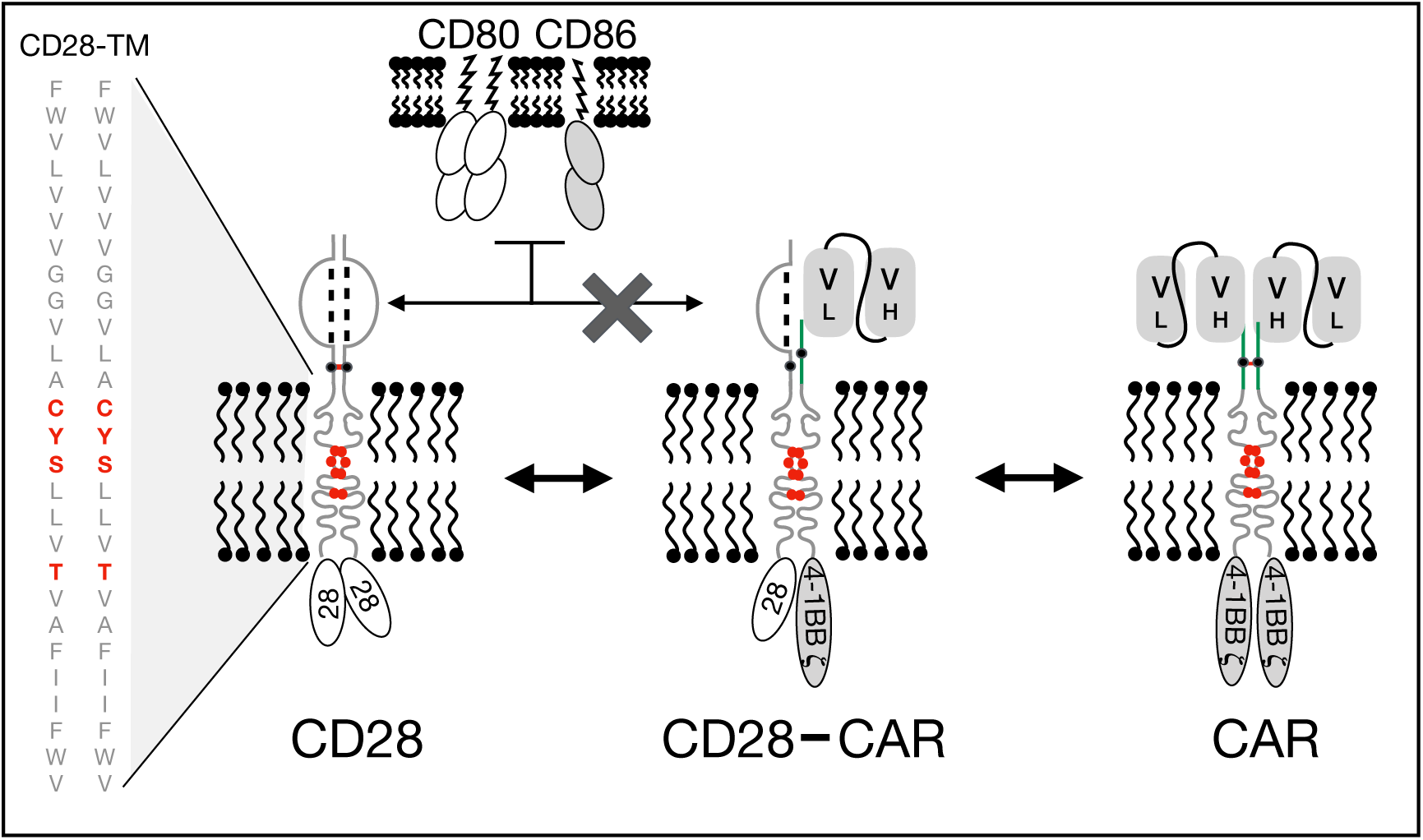

## Introduction

Chimeric antigen receptor (CAR)-engineered T cells are emerging as promising therapies for otherwise untreatable diseases (1). The FDA has approved two anti-CD19 CAR (CD19-CAR) T-cell products, tisagenlecleucel (CTL-019, KYMRIAH, Novartis Pharmaceuticals Corp.) and axicabtagene ciloleucel (KTE-19, YESCARTA, Kite Pharma, Inc.), for the treatment of acute lymphocytic leukemia and relapsed/refractory large B-cell lymphoma. A third CAR-T product, Lisocabtagene Maraleucel (JCAR-17, LISO-CEL, Bristol-Myers Squibb), is currently under review by the FDA for adults with relapsed/refractory large B-cell lymphoma. The success of these CAR-T products can be attributed to their antigen specificity, all conferred by the single chain variable fragment (scFv) of the anti-CD19 antibody clone FMC63, and their intracellular signaling domains (ICD), 28**ζ** for KTE-19 and 4-1BB**ζ** for CTL-019 and JCAR-17. It is worth noting that these products differ in their hinge domain (HD) and transmembrane domain (TMD), CD28-HD/TMD for KTE-19, CD8-HD/TMD for CTL-019, and IgG_4_-HD/CD28-TMD for JCAR-17 (2–4).

In earlier iterations of CAR designs, CD28- and CD8-TMDs were chosen because they are considered to be inert when compared to the CD3ζ-derived TMD that mediated association of the CAR with the endogenous T cell receptor (TCR)/CD3 complexes (5). Emerging evidence, however, suggests potential contributions of the HD and TMD to the function of CAR-T cells. The group of Dr. June first observed unexpectedly sustained proliferation after a single *in-vitro* stimulation of CD28-HD/TMD but not CD8-HD/TMD-based-CAR T cells directed against mesothelin (6). More recently, the group of Dr. Mackall demonstrated that replacing a CD8-HD/TMD with a CD28-HD/TMD lowers the threshold for CAR activation to CD19 in an ICD-independent fashion (7). These results corroborate the finding reported by the group of Dr. Kochenderfer showing that T cells with CD28-HD/TMD-containing CARs secrete higher levels of interferon-γ upon CAR stimulation (8, 9). In a recent clinical trial, the Kochenderfer’s group observed lower incidences of neurotoxicity with a new humanized CD19-CAR-T cell product engineered with CD8-HD/TMD when compared to KTE-19 and attributed this difference to the HD/TMD in the two CARs (10).

The mechanisms underlying the differences between CD8-HD/TMD and CD28-HD/TMD domains remain to be defined (11). In the present work, we investigated the impact of CD28-TMD on CD19-CARs in human T cells and made the surprising discovery that CD28-TMD mediates a heterodimeric association of the CAR with the endogenous CD28 receptor.

## Results and discussion

To investigate the role of CAR TMD, we first generated a panel of CD19-CARs differing only by their HD (CD8, CD28 or IgG_4_) and TMD (CD8 versus CD28), all of which have been used to engineer CAR T cells for clinical applications (Figure 1A, Supplementary Figure 1). Each CAR was designed with a MYC tag on the N-terminus of the scFv and a mCherry reporter (Figure 1A). We selected 4-1BB’s ICD to avoid potential interactions with the endogenous CD28. Furthermore, we disrupted the TCR beta chain constant region (*TRBC*) locus using CRISPR/Cas9 to prevent any potential confounding influence by the endogenous TCR (Figure 1B). The *TRBC* gene-disrupted human T cells retained cell surface expression of TCR/CD3 proteins for a few days after editing and could thus be activated with anti-CD3/CD28 beads. Edited CD4^+^ T cells were transduced with various lentiviral CAR constructs by spinoculation two days after activation. On day 9 after stimulation, 87-98% of the cells were CD3-negative, demonstrating successful TCR deletion in the majority of the cells (Figure 1C). Comparable transduction efficiencies were observed across the different CAR constructs, as assessed by mCherry expression and all CAR T cells responded to CD19 re-stimulation (Supplementary Figure 2A-C).

**Figure 1.**
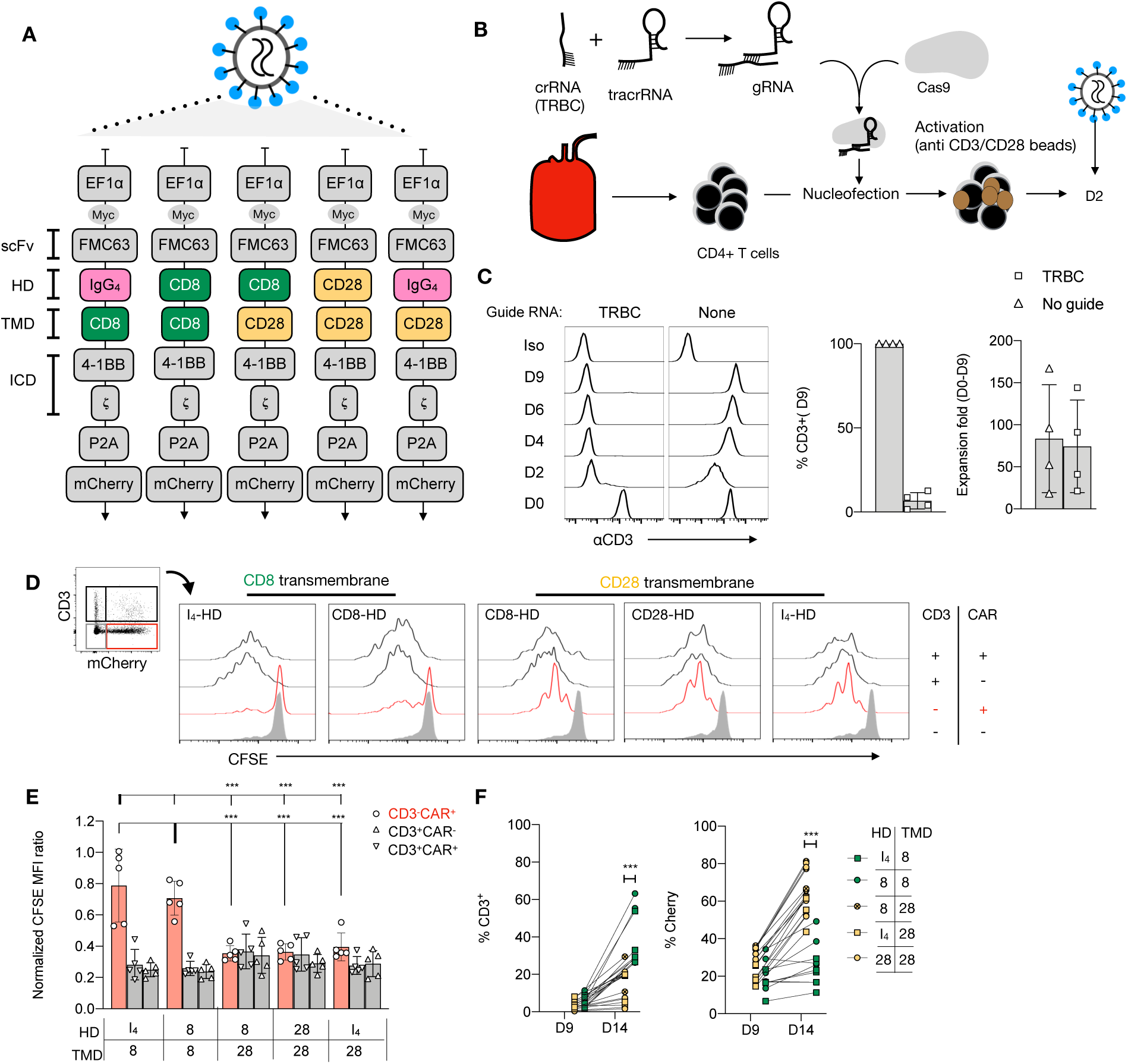
Anti-CD28 stimulation of CD19-CAR T cells is TMD dependent. **(A)** Designs of five chimeric antigen receptors (CAR) against CD19 bearing a 4-1BB co-stimulatory domain and differing by their hinge and transmembrane domain. (**B**) FACS sorted CD4^+^CD127^+^CD25^low^ T cells were electroporated with a CRISPR-Cas9 ribonucleoprotein complex (RNP) targeting the constant region of the TCR β chain gene (*TRBC*), followed by stimulation with anti-CD3/CD28 beads (1:1 ratio). (**C**) Representative results of flow cytometric analysis of CD3 expression over time of cells electroporated with or without RNP. Percentages of residual CD3^+^ population and fold-expansion after 9 days of culture of CD4^+^ T cells electroporated with or without RNPs targeting *TRBC* are shown. Results from 4 independent experiments. (**D**) A representative example of CFSE dilution of a mixed population of CD3^+/-^CAR^+/-^ T re-stimulated with anti-CD3/28 beads. (**E**) Normalized CFSE MFI ratio for CD3^-^mCherry^+^, CD3^+^mCherry^-^ and CD3^+^mCherry^+^ cells was calculated by dividing CFSE MFI of these populations with the MFI of the CD3^-^mCherry^-^ cells in the same culture. Two-way ANOVA was used for statistical analysis (bold line set as reference). (**F**) Percentages of CD3^+^ and mCherry^+^ cells before and 5 days after re-stimulation of edited T cells with anti-CD3/CD28 beads. Unpaired t-test was performed comparing CD8-TMD and CD28-TMD-containing CARs on D14. For **E** and **F**, results shown are a summary of 2 independent experiments using T cells from 5 unrelated donors for each construct. *** p<0.001

Re-stimulation of TCR-edited CAR-transduced T cells, containing a mixed population of CD3^+/-^ and CAR^+/-^ cells, with anti-CD3/CD28 beads on day 9, resulted in the expansion of CD3^+^ T cells that escaped TCR deletion (Figure 1D and E). Surprisingly, TCR-deficient, CD3^-^ CAR^+^ T cells with CARs containing a CD28-TMD, but not CD8-TMD, also proliferated. Consequently, CAR^+^ T cells transduced with CARs containing a CD28-TMD, but not CD8-TMD, were enriched at the end of the 5-day re-stimulation (Figure 1F). The lack of proliferation of CD8-TMD-containing CAR T cells suggests that expansion was not a consequence of bystander effects, such as IL-2 production by the CD3^+^CAR^+^ T cells in the same culture. To determine if this is unique to CARs with 4-1BB-ICD, we repeated the experiment using CARs with a CD28-ICD and observed a similar pattern of proliferation and enrichment of CD3^-^CAR^+^ T cells after anti-CD3/28 bead re-stimulation (Supplementary Figure 3A-D).

To verify that the endogenous CD28 receptor was required for proliferation in response to anti-CD3/CD28 beads, both the *CD28* and *TRBC* genes were deleted in T cells before activation and lentiviral CAR transduction (Figure 2A). CD3^-^CAR^+^CD28^+^ T cells expressing CARs containing a CD28-TMD, but not a CD8-TMD, proliferated in response to anti-CD3/28 beads. Deletion of *CD28* abrogated the ability of CD28-TMD-containing CAR T cells to proliferate in response to anti-CD3/CD28 beads, demonstrating that CD28-mediated activation depends on endogenous CD28 expression (Figure 2A).

**Figure 2:**
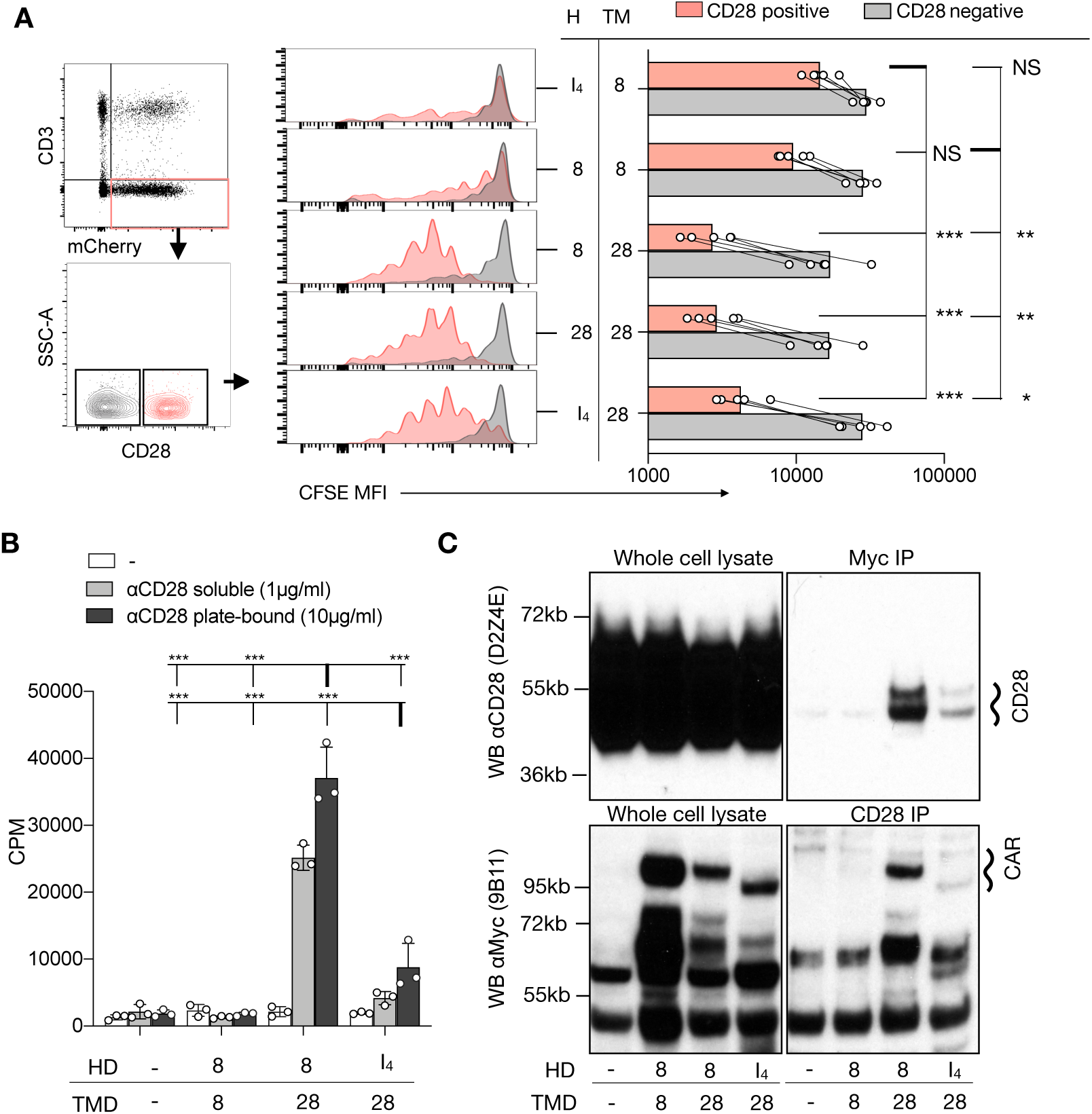
CD28-TMD-containing CARs interact with CD28. **(A)** A mixture of CFSE-labeled CD4^+^ T cells with or without CD3, CD28, and CAR expression. CFSE MFI of 5 independent donors in 2 independent experiments is reported. One-way ANOVA was used for statistical analysis. (**B**) Proliferation of purified CD3^-^CAR^+^ CD4^+^ T cells in response to plate-bound or soluble anti-CD28 stimulation. Results are representative of 3 independent experiments. Two-way ANOVA was used for statistical analysis. **(C)** CD28 or the Myc-tag of CD3^-^CAR^+^ T cells were immunoprecipitated. Western blot analysis of the input (5% of the whole cell lysate) as well as of the precipitated was performed using anti-CD28 (clone D2Z4E) and anti-Myc (clone 9B11). Results are representative of 2-3 independent experiments for each condition. ** p<0.01***, p<0.001 Counts per minute (CPM). Not Statistically Significant (NS).

To determine if CD3^-^ T cells with a CD28-TMD-containing CAR can respond to anti-CD28 stimulation alone without influence from other cells in the culture, we FACS-purified CD3^-^ CAR^+^ cells before re-stimulation with plate-bound or soluble anti-CD28 antibodies (clone CD28.2). For these experiments, we excluded CD28-HD containing CARs to avoid potential interaction mediated by the CD28-HD. The results confirmed that CAR T cells engineered with a CD28-TMD, but not a CD8-TMD, proliferated in response to anti-CD28 alone (Figure 2B). The proliferative response induced by anti-CD28 alone in CD3^-^CAR^+^ T cells was similar in CAR T cells with a 4-1BB or a CD28 costimulatory domain in their ICD (Supplementary Figure 4A). Moreover, anti-CD28 induced secretion of multiple cytokines by CD3^-^CAR^+^ T cells, but not by CD3^+^CAR^-^ or CD3^-^CAR^-^ control cells (Supplementary Figure 4B). Collectively, these results show that CD28-TMD containing CARs can be activated by anti-CD28 without antigen recognition by the CAR or TCR. These results, together with recent reports of phosphorylation of endogenous CD28 upon CAR stimulation (with a CD28-TMD domain), suggest interactions between CD28 and CD28-TMD-containing CARs (12, 13).

To directly determine if CD28-TMD-containing CAR and CD28 physically interact, we performed co-immunoprecipitation experiments. CD28-TMD-containing, but not CD8-TMD-containing, CARs co-immunoprecipitated with endogenous CD28. Conversely, endogenous CD28 co-immunoprecipitated with CD28-TMD-containing, but not CD8-TMD-containing, CARs, demonstrating that the CD28-TMD of the CAR interacted with the endogenous CD28 receptor (Figure 2C). It is worth noting that CD8-HD/CD28-TMD CARs and CD28 co-immunoprecipitated more efficiently when compared to the IgG_4_-HD-CD28-TMD construct, which is consistent with the better proliferation observed with CD8-HD/CD28-TMD CAR upon anti-CD28 stimulation (Figure 2B). We speculate that this difference may be due to the very short IgG_4_-HD (Supplementary Figure 1) which may not be as flexible as other hinges, leading to steric hindrance by the globular scFv domain (14). Alternatively, the membrane proximity of the cysteine in the IgG_4_-HD may not readily form disulfide bonds with the cysteine in the CD28 hinge of the endogenous CD28 receptor.

Next, we generated a series of CD28-TMD CAR mutants to determine the molecular basis of the CAR-CD28 interaction. We first mutated the two glycine G160L and G161L (M1) that may function as part of a glycine-zipper motif, a process known to control TMD dimerization (15). The second mutation replaced the C165 cysteine to alanine, as cysteine can form disulfide bonds (M2). The third (M3) and fourth (M4) sets of mutations were made on amino acids targeting either the two bulky hydrophobic tryptophans at the border of the TMD (W154L and W179L) or the four amino-acid residues in the core of the TMD (C165L, Y166L, S167L, and T171L), as cysteine could form a disulfide bond and others may hydrogen bond across the interface (Figure 3A). All CARs with TMD-mutants were readily expressed on the cell surface (Figure 3A). The various CD3^-^CAR^+^ cells with mutated CD28-TMD were examined for their ability to proliferate to anti-CD28 stimulation. Based on the level of mCherry expression, CD3^-^ CAR^+^ cells were defined as low, intermediate, or high CAR expressors. CAR T cells with the wild type CD28-TMD (CD28^WT^-TMD) proliferated to anti-CD3/CD28 stimulation regardless of the level of CAR expression (Figure 3B-C). The CD28^M4^-TMD, but not the other TMD-mutants, abrogated the proliferation of CD3^-^CAR^low^ cells and significantly reduced proliferation of CD3^-^CAR^int^ cells with either CD8-HD or IgG_4_-HD (Figure 3B-C). Interestingly, CD3^-^CAR^high^ T cell proliferation was only weakly affected by M4 mutations. The proliferation of CD3^-^CAR^high^ T cells was not observed under conditions when the cells were not re-stimulated, demonstrating that activation was dependent on anti-CD28 stimulation, and not a result of autonomous CAR tonic signaling (Figure 3C). It has been reported that CD28-mediated co-stimulation can induce coalescence of membrane microdomains enriched for signaling molecules resulting in enhanced T cell activation (16). Thus, cells expressing high levels of CAR may become sufficiently activated to proliferate by CD28-induced membrane compartmentalization.

**Figure 3:**
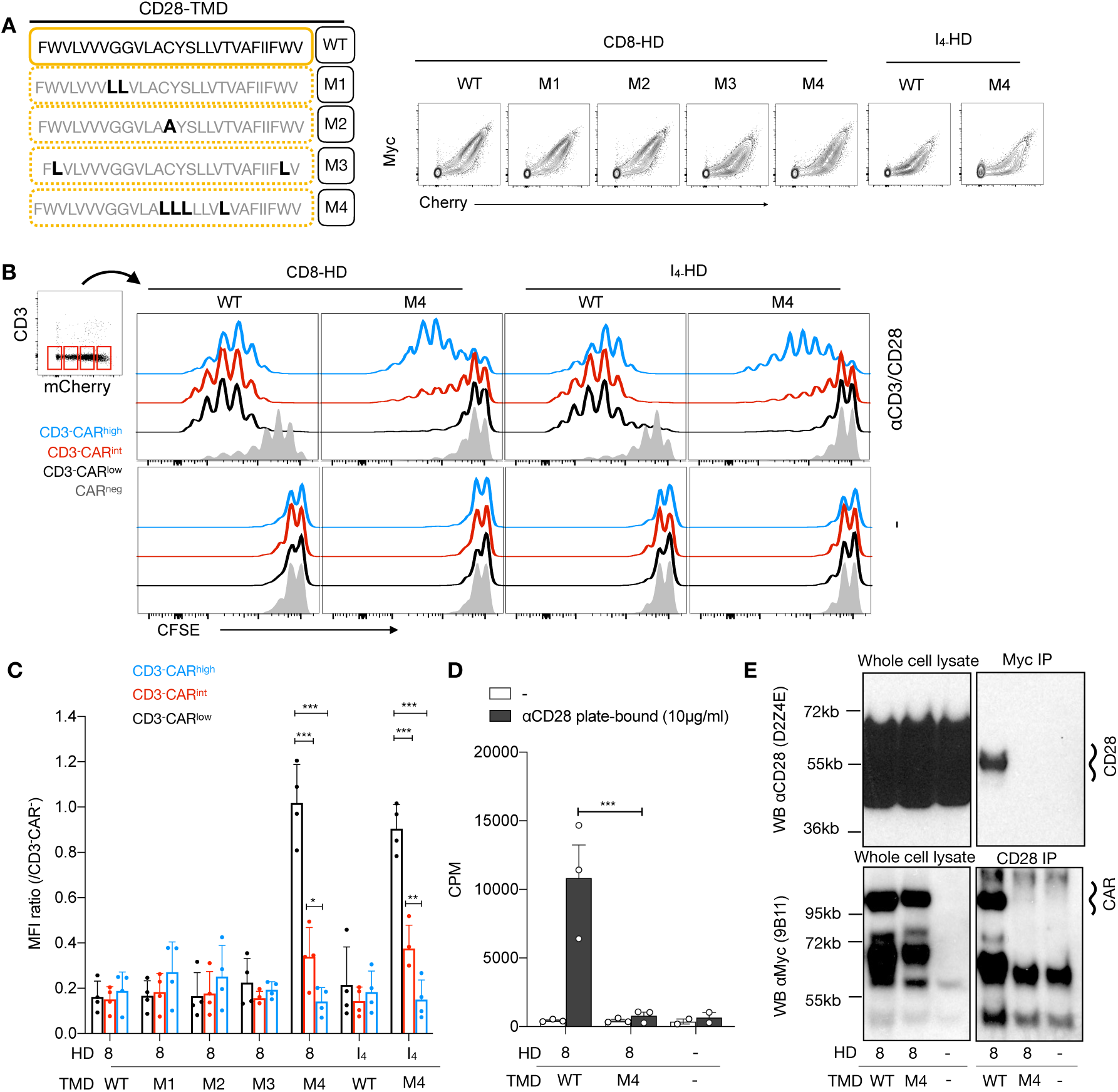
The dimerization of the CD28-TMD depends on a core of four amino acids. (**A**) Diagram representing the amino acid sequence of the wild type and four mutants of the CD28-TMD. A representative example of MYC and mCherry expression for each mutant is shown. (**B**) A representative example of CFSE dilution of a mixed population of CD3^+/-^CAR^+/-^ T cells re-stimulated with anti-CD3/CD28 beads. (**C**) Normalized CFSE MFI ratio for CD3-CAR^low/int/high^ was calculated by dividing the MFI of each of these population with the MFI of CD3^-^mCherry^-^ cells within the same culture. A summary of results using T cells from 4 unrelated donors in 2 independent experiments is shown. (**D**) Proliferation of purified CD3^-^ CAR^+^ T cells in response to plate-bound anti-CD28 stimulation. Results represent the mean of 3 independent experiments. (**E**) CD28 or the Myc-tag of CD3^-^CAR^+^ T cells were immunoprecipitated. Western blot analysis of the input (5% of the whole cell lysate) as well as of the precipitated was performed using anti-CD28 (clone D2Z4E) and anti-Myc (clone 9B11). Results are representative of 2 independent experiments for each condition. Two-way ANOVA were used for statistical analysis. * p<0.05, ** p<0.01***, p<0.001

To confirm that the CD28^M4^-TMD mutant disrupted the interaction between CD28 and the CAR, we sorted CAR T-cells engineered either with a CD8-HD/CD28^WT^-TMD, or with CD8-HD/CD28^M4^-TMD and re-challenged them with plate-bound anti-CD28 (Figure 3D). In this assay, only CAR T cells with a CD28^WT^-TMD showed significant proliferation, as measured by radiolabeled-thymidine incorporation. Importantly, co-immunoprecipitation of the endogenous CD28 and CD28-TMD-containing CAR was abrogated by the M4-mutant, demonstrating that the four amino acids in the core of the CD28-TMD are necessary for CAR-CD28 heterodimerization (Figure 3E). Our finding is consistent with a recent report of the existence of an evolutionarily conserved YxxxxT motif in the CD28TMD that mediates CD28 homodimerization (23). This study further demonstrates that this motif also mediates heterodimerization between synthetic CAR and the endogenous CD28.

To determine if the natural ligands of CD28, CD80 and CD86, can activate CARs by engaging CD28-CAR heterodimers, we stimulated CAR T cells with different HD and TMD with CD19-deficient Raji cells that express high levels of CD80 and CD86 (Supplementary Figure 5A). CD19^null^ Raji induced CAR T cell activation, although less than that induced by the CD19^+^ Raji cells (Supplementary Figure 5B, Figure 4A). This “off-target” activation was mostly seen in T cells with high CAR expression (Figure 4A-B). Moreover, CAR T activation by CD19^null^ Raji was significantly reduced by CTLA-4 Ig, a high-affinity competitive inhibitor of CD28 by binding to CD80 and CD86 (Figure 4A-B), demonstrating that the off-target activation of CAR T cells is predominantly driven by the CD28 interaction with CD80 and CD86. CD28^M4^-TMD did not markedly affect the off-target activation of the IgG_4_-HD CAR, likely because of the weak heterodimerization between IgG_4_-HD/CD28^WT^-TMD CAR and CD28 (Figure 2C). Intriguingly, the CD28^M4^-TMD significantly potentiated off-target activation of CD8-HD containing CAR T cells (Figure 4B). CD80 and CD86 are reported to engage CD28 homodimers (17) and our results suggest they are unable to recognize CD28 in CD28-CAR heterodimers. This further suggests that the off-target CAR T cell activation by CD80/86 was mediated by the endogenous CD28 homodimers, which may induce CAR clustering through membrane compartmentalization and signaling especially in cells with high CAR expression (Figure 3C).

**Figure 4:**
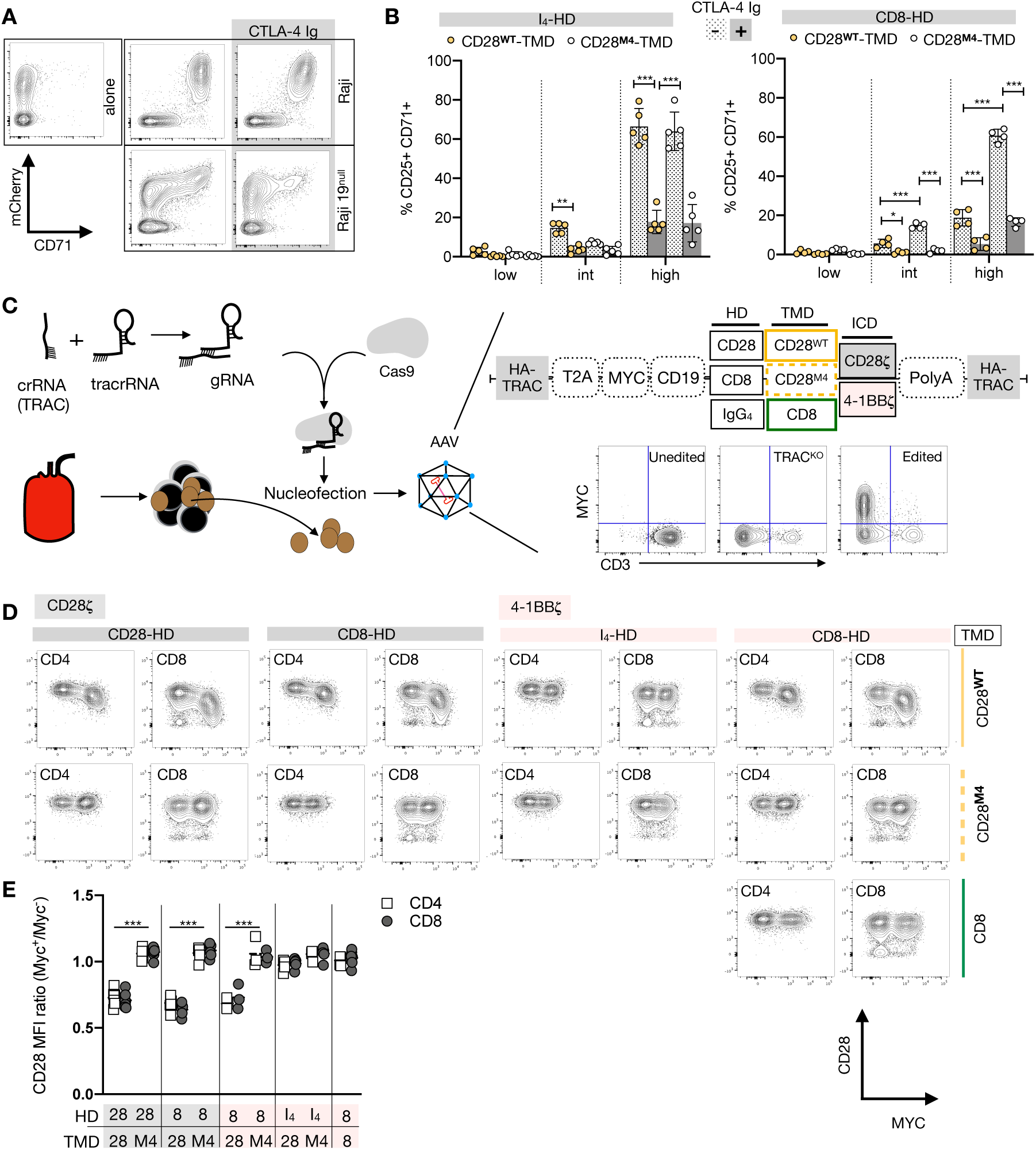
CAR-CD28 heterodimers are B7-unresponsive but reduce CD28 expression. (**A**) Representative example showing CD71 upregulation in CAR T cells containing an IgG_4_-HD/CD28-TMD co-cultured for 48h with irradiated (4000Rad) CD19-wild type or deficient Raji cells with or without CTLA-4 Ig. (**B**) CD25^+^CD71^+^ T cells were analyzed in low, intermediate (int) or high mCherry-expressing CAR-T cells using the gating strategy described in Supplementary Figure 5. Data were pooled from 4 independent experiments using T cells from 4-5 unrelated donors. (**C**) Editing strategy and homology-directed repair-mediated integration into the *TRAC* locus of various CD19 CARs using an AAV-6 transduction protocol. (**D**) Myc and CD28 expression in a representative example analyzed 6 days after editing and beads removal. (**E**) CD28 MFI ratio was calculated by dividing CD28 MFI of Myc^+^ cells by Myc^-^ cells in the same culture. Pooled data from 3-4 independent experiments across 5 unrelated donors are shown. Each dot represents one independent editing condition. Two-way ANOVA was used for statistical analysis. *p<0.05, *** p<0.001.

A previous report has shown that the percentages of CD28^+^ CD19-CAR T cells engineered with a CD28-TMD are significantly lower when compared to CD19-CAR T cells engineered with a CD8-TMD. This correlated with a significantly lower number of CD28-TMD-containing CAR T cells in the peripheral blood one month after infusion into patients, suggesting that the reduced CD28 expression might have impaired CAR T cell persistence (10). We thus examined if CAR-CD28 heterodimerization regulates CD28 expression. Since lentiviral transduction resulted in a wide range of CAR expression levels that influenced on- and off-target T cell activation, we expressed various CARs by knocking them into the TCR alpha constant (*TRAC*) gene locus using homology-directed repair that provided more homogenous expression of the CAR (Figure 4C) (18). Knock-in efficiencies ranged between 17-72% across the various CAR constructs but the levels of CAR expression were similar regardless of the differences in editing efficiency (Supplementary Figure 6A). In addition, all CAR T cells proliferated comparably upon stimulation with CD19^+^ NALM-6 target cells (Supplementary Figure 6B), demonstrating M4 mutations in the CD28-TMD did not impact on-target CAR activation. Six days after CAR knock-in prior to exposing the cells to target cells, we observed a 26-51% reduction in CD28 staining on CAR T cells containing a CD28^WT^-TMD as compared to a CD28^M4^-TMD with either a CD8-HD or CD28-HD (Figure 4D-E). This reduction was seen in both CD4 and CD8 T cells with CARs containing either 28**ζ** or 4-1BB**ζ** ICD. The downregulation of CD28 in CAR T cells was only minimal with CAR T cells engineered with an IgG_4_-HD/CD28^WT^-TMD, echoing the earlier result of inefficient CAR-CD28 heterodimerization in the context of IgG_4_-HD. However, the reduced staining is not likely due to the inability of the anti-CD28 antibody to bind to CD28 in heterodimers since we could activate heterodimer-expressing CAR T cells with anti-CD28 (Figures 2 and 3). Rather, reduced CD28 staining most likely reflected a loss of cell surface expression of CD28 protein. CD28 downmodulation occurred days after CAR engineering, without exposure to target antigen or CD28 ligands, suggesting that this might be mediated at the level of protein synthesis and stability. A recent report showed that CD28 cell-surface expression depended on the dimerization of the CD28 TMD (23). Thus, the reduction of CD28 expression likely resulted from reduced homodimerization of endogenous CD28 due to the recruitment of CD28 into CAR heterodimers. The finding of CAR CD28-TMD-dependent downmodulation of CD28 may also explained our observation that CD80/86-mediated off-target activation was less pronounced with the dimerization-competent CD28-TMD CAR than with the dimerization-disabled CD28-TMD^M4^ CAR (Figure 4B).

In conclusion, we have discovered that the CD28-TMD mediates CAR and CD28 heterodimerization via a core of up to four polar amino acids, of which the tyrosine (Y), threonine (T), and serine (S) can form hydrogen bonds (19). In fact, the CAR-CD28 heterodimerization may be mediated by the YxxxxT motif in the CD28-TMD solely, as recently demonstrated in the context of CD28 dimerization (23). Importantly, the efficiency of the heterodimerization is modulated by the HD of the CAR. The CAR-CD28 heterodimer mediates CAR T-cell activation after anti-CD28 stimulation independently of TCR and CAR cognate antigens but did not respond to natural CD28 ligands, CD80 and CD86. Yet, CAR-CD28 heterodimerization is associated with reduced cell surface expression of CD28. Together, these results demonstrate an active role of the TMD and the HD on CAR T cells. Future investigations are needed to understand the contribution of the CAR-CD28 heterodimerization to CAR T cell activation, survival, exhaustion, and CAR T-cell associated toxicities. Thus, our findings suggest that CAR TMDs may modulate CAR T activities by association with their endogenous partners. Optimization of CAR designs should incorporate consideration of TMD-mediated receptor interactions.

## Materials and methods

### Human T cells isolation

Human T cells were obtained by purchasing whole blood units from STEMCELL technologies (Vancouver, Canada). STEMCELL reported that these products are distributed using Institutional Review Board (IRB)-approved consent forms and protocols. Peripheral blood mononuclear cells were isolated by Ficoll density gradient centrifugation and T cells were further enriched using the EasySep Human T Cell Isolation Kit (STEMCELL) per manufacturer’s instructions. Enriched T cells were stained with antibodies against CD4, CD25, and CD127 and CD4^+^CD127^+^CD25^low^ conventional T cells were purified by fluorescence-activated cell sorting (FACS). Alternatively, enriched T cells were directly edited and activated with anti-CD3/CD28 beads and IL-2 (300 IU/mL, Prometheus laboratories, Nestle Health Science, Lausanne Switzerland). Cells were either used fresh or cryopreserved in fetal calf serum (FCS) with 10% DMSO and used later after thawing. When frozen cells were used, cells were thawed and cultured overnight in 300 IU/mL recombinant human IL-2 before editing and cell activation.

### Genome editing using ribonucleoprotein complex

Ribonucleoprotein complexes were made by complexing CRISPR RNAs (crRNAs) and trans-activating crRNAs (tracrRNA) chemically synthesized (Integrated DNA Technologies (IDT), Coralville, IA) with recombinant Cas9 protein (QB3 Macrolab, UC Berkley, CA) as previously described (20). Guide RNA sequences used for gene editing were:

1. T cell receptor beta chain constant region (TRBC): CCCACCAGCTCAGCTCCACG;
2. CD19: CGAGGAACCTCTAGTGGTGA;
3. CD28: TTCAGGTTTACTCAAAAACG.

Lyophilized RNAs were resuspended at 160µM in 10mM Tris-HCl with 150mM KCl and stored in aliquots at −80°C. The day of electroporation, crRNA and tracrRNA aliquots were thawed and mixed at a 1:1 volume and annealed for 30 minutes at 37°C. The 80 µM guide RNA complex was mixed at 37°C with Cas9 NLS at a 2:1 gRNA to Cas9 molar ratio for another 15 minutes. The resulting ribonucleoprotein complex (RNP) was used for genome editing. To delete TCR or CD28 genes, 1 x10^6^ T cells were mixed with appropriate RNP and electroporated using a Lonza 4D 96-well electroporation system (pulse code EH115). For generating CD19^-^ variants of Raji cells, parent Raji cells (ATCC^®^ CCL-86™, Manassas, VA) were electroporated (pulse code EH140) with ribonucleoprotein complex targeting CD19 and the CD19-negative fraction was purified by FACS.

### Activation and lentiviral transduction of CD4^+^ T cells

CD4^+^ T cells were electroporated prior to stimulation with anti-CD3/CD28 beads (Dynabeads Human T-Activator CD3/CD28, Thermo Fisher Scientific (ThermoFisher), Waltham, MA). Cells were cultured in RPMI supplemented with 10% FCS and 300 IU/mL of IL-2 for the first two days after electroporation and reduced to 30 IU/mL of IL-2 for CD4 T cells and 100 IU/mL for bulk T cells thereafter. Lentiviruses encoding CD19-I_4_-28-4-1BB**ζ**-T2A-EGFRt and CD19-I_4_-28-28**ζ**-T2A-EGFRt were provided by Juno Therapeutics (Bristol-Myers Squibb, New York, NY). Other lentiviral constructs, depicted in Figure 2A, were cloned into the pCDH-EF1-FHC vector (Addgene plasmid #64874, Watertown, MA) as previously described (21). Briefly, genes encoding CAR constructs were purchased as gblocks (IDT) (21, 22) and amplified by PCR and cloned into the pCDH vector using Infusion cloning tools (Takara Bio, Kusatsu, Japan). Sequences for all clones used in subsequent experiments were confirmed by sequencing. All pLX302-based lentiviruses were produced and tittered by the viral core of UCSF. All lentiviruses were aliquoted and stored at −80°C until use. Transduction was performed on day 2 after CD4^+^ T cell activation at a multiplicity of infection of 1 by spinoculation (1200 g, 30 minutes, 30°C) in medium supplemented with 10% FCS and 0.1 mg/mL of protamine. Regarding AAV production, helper plasmid 30 μg of pDGM6 (a kind gift from YY Chen, University of California, Los Angeles), 40 μg of pAAV helper, and 15 nmol PEI were utilized. AAV6 vector production was carried out by iodixanol gradient purification. After ultracentrifugation, the AAVs were extracted by puncture and further concentrated using 50 mL Amicon column (Millipore Sigma Burlington, MA) and directly titrated on primary human T cells. Transduction was performed on day 2 after T cell activation.

### *In vitro* activation of gene-edited CAR T cells

For some experiments, day-9 cells were re-stimulated without separating edited and transduced cells. For proliferation assays, the cell mixtures were stained with 2.5 µM carboxyfluorescein diacetate succinimidyl ester (CFDA SE, ThermoFisher, referred to as CFSE) before restimulation with anti-CD3/CD28 beads. For some other experiments, day-9 cells were separated by FACS to purify CD3^+^ and CD3^-^ T cells with or without CAR. For assessing early T cell activation, purified CAR T cells were stimulated with soluble anti-CD28 (clone CD28.2, 1 µg/mL, BD Pharmigen), parental CD19^+^ Raji cells, or CD19^deficient^ Raji cells for 2 days. In some cultures, CTLA-4 Ig (kindly provided by Dr. Vincenti, UCSF) was added at a concentration of 13.5 µg/mL. For measurements of proliferation, purified cells were stimulated with soluble anti-CD28 (clone CD28.2, 1 µg/mL, BD Pharmigen), plate-bound anti-CD28 (clone CD28.2, 10 µg/mL), or soluble anti-CD3 (clone HIT3α 2 µg/mL. BD Pharmigen). After 48 hours, a portion of the supernatant was collected and analyzed for cytokine secretion using multiplexed Luminex (Eve Technologies, Calgary, Canada). The cells were then pulsed with 0.5µCi of ^3^H thymidine and cultured for another 16-18 hrs before harvesting the cells to determine the level of ^3^H thymidine incorporation using a scintillation counter.

### Flow cytometry

The following antibodies were used for phenotyping and proliferation assays: anti-CD3-PE/Cy7 (clone SK7, Biolegend, San Diego, CA), anti-CD4-PerCP (clone SK3, BD Pharmigen, San Jose, CA), anti-CD4 A700 (clone RPA T4, Biolegend), anti-CD19 APC (clone HIB19, BD Pharmigen), anti-CD25 APC (clone 2A3, BD Pharmigen), anti-CD71 FITC (clone CY1G4, Biolegend), anti Myc FITC or APC (clone 9B11, Cell Signaling, Danvers, MA), anti-FMC19 idiotype APC (Juno therapeutics), anti-EGFRt PE (Juno therapeutics), anti-CD28 APC (clone 28.2, Biolegend), CD8 APC-Cy7 (clone SK1 Biolegend). DAPI (ThermoFisher, Waltham, MA) was used to stain dead cells for exclusion during analysis. For *in vitro* mixed lymphocyte reactions or CFSE *in vivo* analysis, Fc-block (Sigma-Aldrich, St. Louis, MO) was used (20 µg/mL) 5 minutes prior to surface staining. Flow cytometric analyses were performed on an LSRII flow cytometer (BD Pharmigen). Fluorescence-activated cell sorting was performed on FACSAria III (BD Pharmigen). All flow cytometry data were analyzed using Flowjo software (Tree Star, Ashland, OR).

### Immunoprecipitation

FACS-purified CD3^-^CAR^+^ or CD3^-^CAR^-^ CD4^+^ T cells (8 x 10^6^ each) were lysed in Pierce™ IP Lysis Buffer (ThermoFisher) supplemented with Complete Protease Inhibitor Cocktail (Roche, Basel, Switzerland) for 30 minutes using a vertical rotator. Cell lysis was completed by briefly sonicating cells using a Q500 sonicator (QSonica, Newtown, CT). Pierce™ anti-c-Myc magnetic beads (clone 9E10, ThermoFisher) were used for immunoprecipitation of the CAR. Alternatively, rabbit anti-human CD28 (clone D2Z4E, Cell Signaling) followed by anti-rabbit IgG Pierce™ protein A/G magnetic beads (Thermo Fisher) were used for CD28 immunoprecipitation of the cell lysate according to the manufacturer’s instructions.

### Western blot

Equal masses of protein lysate or equal volumes of immunoprecipitation eluents were loaded into NuPAGE 4-12% Bis-Tris gels (ThermoFisher). After electrophoresis, proteins were transferred onto PVDF membranes (ThermoFisher) using an iBlot 2 Dry Blotting system. After blocking with Tris-buffered saline with 0.1% Tween-20 and 5% bovine serum albumin (TBSTB), membranes were stained with primary and secondary antibodies diluted in TBSTB. The following antibodies were used: mouse anti-Myc (clone 9B11, Cell Signaling), rabbit anti-CD28 (clone D2Z4E, Cell Signaling), HRP-conjugated anti-mouse IgG (Cell Signaling) and HRP-conjugated anti-rabbit IgG (Cell Signaling).

## Abbreviations

CAR: Chimeric Antigen Receptor
HD: Hinge domain
ICD: Intracellular signaling domain
scFv: Single chain variable fragment
TMD: Transmembrane domain,

## Authors contributions

Conceptualization: YDM, QT; Formal analysis: YDM; Funding acquisition: QT; Investigation: YDM, DPN, LMRF, CR, ZCW, RBV; Methodology: YDM DPN, TR, PH, JE; Analysis: YDM, DPN; Resources: QT, JAB, AM, JAW. Supervision: QT, JAB. Writing-original draft: YDM, QT. Writing-review & editing: YDM, QT, DPN, LMRF, CR, AM, JE, TR, FVG, ZCW, RBV, JAW, JAB.

## Acknowledgment

YDM is supported by the Swiss National Science Foundation (Advanced Postdoctoral Mobility Grant no. P300PB_174500). LMRF is the Jeffrey G. Klein Family Diabetes Fellow. QT and JAB acknowledge research grants from the NIDDK (UC4 DK116264 and P30 DK063720). DN was supported on a Damon Runyon Fellowship (DRG-2204-14). JAW thanks support from R35GM122451 and the Harry and Dianna Hind Professorship. A.M. holds a Career Award for Medical Scientists from the Burroughs Wellcome Fund, is an investigator at the Chan–Zuckerberg Biohub, a member of the Parker Institute for Cancer Immunotherapy (PICI) and is a recipient of The Cancer Research Institute (CRI) Lloyd J. Old STAR grant. The Marson lab has received funds from the Innovative Genomics Institute (IGI) and the Parker Institute for Cancer Immunotherapy (PICI). We acknowledge Juno therapeutics for providing I_4_-28-EGFRt viruses (with a 4-1BB or CD28 co-stimulatory domain).

